# Speed amplitude and period affect gait variability and step followability under sinusoidal speed changing conditions

**DOI:** 10.1101/2024.10.15.618616

**Authors:** Kiyotaka Motoyama, Takehiro Tashiro, Akira Saito, Masahiro Horiuchi, Taisuke Sakaki, Daijiro Abe

**Author notes:** Corresponding Author: Daijiro Abe, Ph.D., Center for Health and Sports Science, Kyushu Sangyo University, 2-3-1 Matsukadai, Higashi-ku, Fukuoka 813-8503, Japan.

## Abstract

**Background:** The time courses of the joint elevation angles of the thigh, shank, and foot in one stride during walking can be well approximated by a “*plane*” in a triaxial space. This intersegmental coordination (IC) of the lower limb elevation angles is called planar covariation law. Thickness of the IC *plane* is associated with gait variability. This study aimed to examine how anteroposterior and lateral gait variabilities are influenced by sinusoidal speed changes with different amplitudes (±0.33 vs. ±0.67 m·s^-1^) and periods (30 vs. 60 s). We also questioned which limbs are responsible for the step variabilities in these conditions.

**Methods:** Eighteen young adults walked on a treadmill under sinusoidal speed-changing conditions with different amplitudes (±0.33 vs. ±0.67 m·s^-1^) and periods (30-vs. 60-s). Using 3D motion analysis system, we quantified the IC *plane* thickness, coefficient of variance of step width (CV_SW_), time delay of step length (TD_SL_), and step frequency (TD_SF_). We applied 2-way statistical parametric mapping for the time courses of each limb angle during the acceleration and deceleration phases.

**Results:** The IC *plane* thickness was greater in the ±0.67 m·s^-1^ condition than in the ±0.33 m·s^-1^ condition. Periods and amplitudes did not affect CV_SW_, TD_SL_, and TD_SF_. In the middle gait cycle, the thigh and shank were delayed in the greater amplitude condition during the acceleration phase but proceeded in the same condition during the deceleration phase.

**Discussion:** Sinusoidal speed amplitude influenced anteroposterior gait variability, but not lateral gait variability, regardless of period, even in healthy young adults. More distal limbs were delayed in the greater speed amplitude condition during the acceleration phase, whereas more proximal limbs proceeded in that condition during the deceleration phase, indicating that these different behaviors of the lower limb segments could be related to step variabilities.

## Introduction

The trajectory of the elevation angles of the thigh, shank, and foot in a gait cycle can be well approximated by a “*plane*” in a triaxial space (*Lacquaniti, Grasso & Zago 1999*), called the planar covariation law (PCL) (*Hicheur, Terekhov & Berthoz 2006*; *Ivanenko et al., 2008*; *Aprigliano et al., 2017*; *Dewolf et al., 2019*; *Abe et al., 2022*). This approach contributes to showing the lower limb’s spatiotemporal interlimb coordination (IC) during human gait. Moreover, the shape of the IC *plane* was altered by an abrupt perturbation of treadmill speed (*Aprigliano et al., 2017*; *Aprigliano, Monaco & Micera, 2019*). Thus, variability of the planarity of the IC *plane* in a gait cycle may be a result of the responses of individual lower limbs to maintain gait stability against the speed perturbation. Indeed, an increased degree of gait speed perturbations did not modify the planarity of the IC *plane* during compensatory behavior in the unperturbed leg (*Aprigliano et al., 2017*), so that the planarity of the IC *plane* has been considered one of the inherent and robust kinematic universalities across several gait-related motor tasks (*Lacquaniti, Grasso & Zago 1999*; *Ivanenko et al., 2008*). Some previous studies reported that gait speed modified the planarity of the IC plane (*Ivanenko et al., 2008*; *Chiu & Chou, 2012*; *Ogaya et al., 2016*; *Dewolf et al., 2018, 2019*), whereas others did not (*Bleyenheuft & Detrembleur, 2012*; *Krasovsky et al., 2014*; *Wallard et al., 2018*). Thus, whether the planarity of the IC *plane* is dependent on the gait speed remains controversial. Notably, these previous studies tested the planarity of the IC *plane* at several steady-state gait speeds ranging from 1-7 km·h^-1^ in various age groups and patients with pathological gait, indicating that other factors rather than gait speed itself may influence gait variability.

In our daily lives, passive gait speed changes necessarily occur based on changes in surface conditions (*Gast, Kram & Riemer, 2019*) and visual illusion (*O’Connor & Donelan, 2012*). A sinusoidal speed-changing protocol is particularly available to evaluate gait variability under speed-changing conditions for several reasons. First, it can involve a wide range of gait speed (*Abe et al., 2022, 2023*; *Horiuchi et al., 2023*). Second, a consecutive spatiotemporal adjustment of the lower limbs is required for walkers without an abrupt perturbation (*Abe et al., 2022, 2023*; *Horiuchi et al., 2023*). Third, the PCL concept can be established regardless of gait speed (*Ivanenko et al., 2008*; *Bleyenheuft & Detrembleur, 2012*; *Dewolf et al., 2019*). Accordingly, we have recently examined the effects of sinusoidal periods of 30-, 60-, and 120-s with a ±0.56 m·s^-1^ (±2 km·h^-1^) amplitude on gait variability and found that the planarity of the IC *plane* was not significantly different among these three periods (*Abe et al., 2022*). However, no previous studies have investigated how the magnitude (amplitude) of gait speed changes in a sinusoidal manner influences the planarity of the IC *plane*. Although a limited number of previous studies have examined the planarity of the IC *plane* under speed-changing conditions (*Aprigliano et al., 2017*; *Aprigliano, Monaco & Micera, 2019*; *Abe et al., 2022, 2023*), the IC *plane* thickness varied with gait speed (*Ivanenko et al., 2008*; *Chiu & Chou, 2012*; *Ogaya et al., 2016*; *Dewolf et al., 2018, 2019*). Thus, these previous results provide a hypothesis that the greater the magnitude of sinusoidal gait speed change, the greater the variability of the IC *plane* planarity. A stable gait with controlled multiple joints must be maintained by continuous adjustments of stride length (SL) and step frequency (SF), so that the time delay (TD) of stride length (TD_SL_) and step frequency (TD_SF_) could reflect delayed adjustment of the lower limbs against sinusoidal speed change (*Abe et al., 2023*; *Horiuchi et al., 2023*). This is because step variabilities refer to the ability of the neuromuscular system to adapt to changing gait conditions (*Chiu & Chou, 2012*; *Ogaya et al., 2016*). In a greater speed amplitude condition, TD_SL_ and TD_SF_ in association with lateral gait variability evaluated by step width (SW) variability (*Brach et al., 2005*; *Kang et al., 2008*; *Skiadopoulos et al., 2020*; *Abe et al., 2022, 2023*; *Horiuchi et al., 2023*) would be greater because the neuromuscular system may not have sufficient time to achieve appropriate adjustment of the lower limbs at a greater amplitude of sinusoidal speed change. Accordingly, it was also hypothesized that the greater the speed amplitude, the larger the TD_SL_, TD_SF_, and SW variabilities. In addition, we further questioned which limb(s) are attributed to a followability of SL-SF combinations against sinusoidal speed change. This study aimed to examine the effects of amplitude (magnitude) and period of sinusoidal speed change on the variabilities of the IC *plane* planarity, SW variability, and followability of SL and SF.

## Materials & Methods

### Participants

This study enrolled 18 healthy young adults (7 men and 11 women; 20.7 ± 1.0 years old, mean ± standard deviation [SD]) without injuries in this study. Their body height and mass were 1.649 ± 0.067 m and 60.9 ± 7.9 kg, respectively. An ethical committee established in Kyushu Sangyo University (No. 2019-0002) approved all procedures. Following the Declaration of Helsinki, all participants signed a written informed consent after being provided information about the purposes, experimental procedures, and possible risks of this study.

### Protocols and motion analysis

We instructed the participants to put on compression shirts, half spats, and the same shoes in different sizes (ADIZERO-RC, Adidas Japan, Tokyo). The participants started walking on a motor-driven treadmill (TKK3080, Takei Scientific Instruments, Niigata, Japan) at 1.33 or 1.25 m·s^−1^ for 30-s for the baseline speed (i.e., midpoint speed during sinusoidal walking), followed by a preliminary habituation and warming-up walk. Subsequently, the treadmill speed was changed in a sinusoidal manner of 60- and 30-s periods with speed amplitudes of ±0.33 m·s^−1^ (±1.2 km·h^−1^) and ±0.67 m·s^−1^ (±2.4 km·h^−1^) in a randomized order (Fig. 1A).

**Figure 1.**
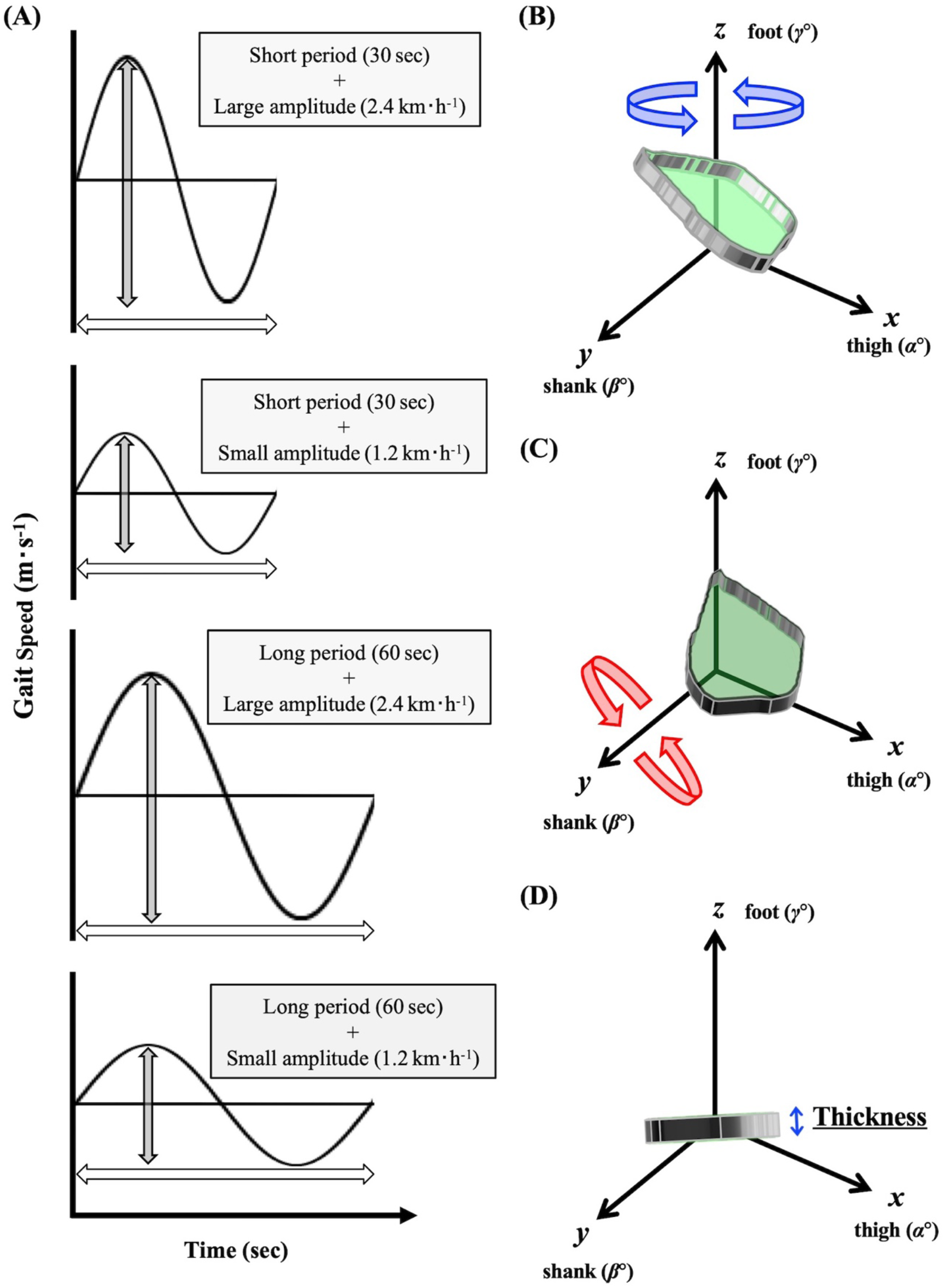
Protocols and Eular’s rotation of planar covariation plane to determine the thickness of interlimb coordination. **(A)** Study protocols. **(B)** The best-fitting loop of the elevation angles of the thigh, shank, and foot is plotted in a squared *x-y-z* space as a “*plane*”. **(C)** The best fitting “*plane*” is rotated around the *z* and *y* axes (shaded in green). **(D)** The *z* angle, at which the smallest standard deviation was obtained, is considered as the thickness of the spatiotemporal interlimb coordination.

Based on our recent studies (*Abe et al., 2022, 2023*), twelve reflective markers were put on both lateral greater trochanters, shoulders (acromion), ankles (lateral malleolus), knees (lateral femur epicondyle), heels (backend of each shoe), and toes (toe of each shoe). Moreover, four additional markers were put on each corner of the treadmill. Motion data were captured using eight high-speed cameras (Kestrel300, MAC3D System, Rohnert Park, CA, USA) with a sampling rate of 100 Hz. The root mean square errors in calculating the three-dimensional (3D) marker locations were within 1.0 mm. The whole gait cycle, defined from the heel-contact to the toe-off, was divided into distinct parts in the range of 0%-100%. We computed the 3 × 3 matrix of the elevation angles of the lower limbs from the marker locations (Fig. 1B) at each time frame. Furthermore, the best-fit 3D covariation loop (IC *plane*) did not perfectly lie on a *plane* (*Hicheur, Terekhov & Berthoz, 2006*; *Bleyenheuft & Detrembleur, 2012*; *Chiu & Chou, 2012*; *Krasovsky et al., 2014*; *Ogaya et al., 2016*; *Wallard et al., 2018*; *Dewolf et al., 2018, 2019*; *Abe et al., 2022, 2023*), and the IC *plane* seems to fluctuate during walking in a sinusoidal speed-changing condition (*Abe et al., 2022, 2023*). Considerably large variations in the IC *plane* thickness could be observed if the shoe sole slightly rubbed the treadmill belt before the real heel strike. Thus, each sinusoidal cycle was continuously repeated thrice to avoid such incomplete motions. Even though the first sinusoidal period was fundamentally used for the subsequent analyses, the second or third cycle was used only when the shoe sole slightly hit the treadmill belt before the real heel strike in the earlier cycles. Accordingly, the largest standard deviation or mean value was not used to represent the IC *plane* thickness, which was considered the smallest standard deviation of the fluctuating IC *plane* in one gait cycle (Fig. 1C).

In a practical computational calculation, the best-fit 3D approximation of the angular covariation is not a dimensionless *plane*. Therefore, based on the definition of Euler’s angle, after detecting the best-fit IC *plane* of the 3D covariation was detected, it was rotated around the *z*-axis (foot elevation angle) as follows (*Abe et al., 2022, 2023*):

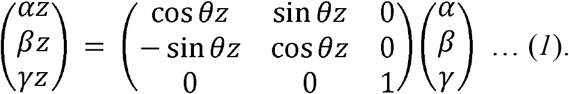

where *α, β*, and *γ* are the original best-fit covariations, and *α*_*z*_, *β*_*z*_, and *γ*_*z*_ are the covariations after rotating around the *z*-axis. The matrix described by Eq. (1) was simultaneously rotated around the *y*-axis (knee elevation angle) as follows:

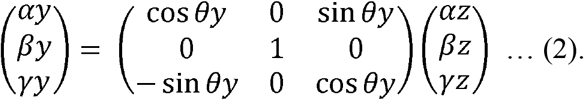

where *α*_y_, *β*_*y*_, and *γ*_*y*_ are the covariations after rotating around the *y*-axis. Thus, the *plane* was rotated by a combination of the matrices 1 and 2 as follows:

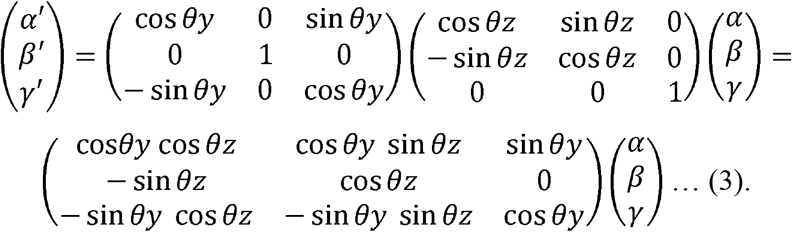

Considering both rotation angles, *θ*_z_ and *θ*_y_, ranging from 0° to 179°, 32,400 (180 × 180) combinations can be defined. Subsequently, the IC *plane* thickness was calculated in a non-arbitrary computational space.

The motion data were also used to calculate the TD_SL_ and TD_SF_ against sinusoidal speed change. The following equation was used to approximate SL and SF:

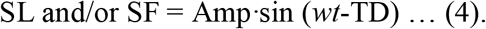

where Amp, *ω*, and *t* represent amplitude, angular frequency, and time (ms), respectively. The SW was quantified as the lateral distance between both heel makers in each step (*Abe et al., 2022, 2023*) because it was reported to be less dependent on the gait speed (*Kang et al., 2008*; *Ciprandi et al., 2018*; *Oliveira et al., 2017*). Thus, the SW was measured during the whole first period (30-or 60-s) to calculate the coefficient of variance (CV_SW_; %) as the SW variability.

### Statistical analysis

The software G*Power 3.1 (*Faul et al., 2007*) was used to estimate the required number of participants with an effect size of 0.25 determined by a partial *η*^2^ of 0.06, which is the medium effect size measure indicating the total variance in testing explained by the within-participants variables (*Cohen, 1988*). At least 16 participants were required in this study after setting a statistical power of 0.8, an alpha level of 0.05 and correlations among repeated measures of 0.8. Two-way repeated measures analysis of variance (ANOVA) within periods (30- and 60-s) and amplitudes (±0.33 and ±0.67 m·s^-1^) was performed on the dependent variables using ANOVA 4 on the web. To examine which limb(s) are attributed to TD_SL_ and/or TD_SF_, we applied two-way statistical parametric mapping (SPM) for the relative time series of each limb (*Pataky, Vanrenterghem & Robinson, 2015*). The time series data were divided into the acceleration and deceleration phases. Statistical significance was set at *p* < 0.05. All data were presented as mean ± SD.

## Results

ANOVA showed a significant amplitude effect in the IC *plane* thickness (*F* = 10.286, *p* = 0.005, *η*^2^ = 0.018; Fig. 2A). A main effect of the sinusoidal period (*F* = 0.011, *p* = 0.919, *η*^2^ = 0.000; Fig. 2A) and interaction effect (*F* = 0.234, *p* = 0.635, *η*^2^ = 0.000; Fig. 2A) were not significant. The CV_SW_ trended to be greater in the ±0.67 m·s^-1^ condition than in the ±0.33 m·s^-1^ condition (*F* = 4.402, *p* = 0.051, *η*^2^ = 0.019; Fig. 2B), but this trend was not observed between the 30- and 60- sec period (*F* = 0.083, *p* = 0.777, *η*^2^ = 0.000; Fig. 2B). The TDSL was not significantly different between periods (*F* = 0.069, *p* = 0.796, *η*^2^ = 0.000; Fig. 3A) and amplitudes (*F* = 0.402, *p* = 0.534, *η*^2^ = 0.000; Fig. 3A). The TDSF was the same as the TDSL between periods (*F* = 0.012, *p* = 0.913, *η*^2^ = 0.000; Fig. 3B) and amplitudes (*F* = 0.657, *p* = 0.429, *η*^2^ = 0.001; Fig. 3B). Consequently, the total TD was not significantly different between periods (*F* = 0.090, *p* = 0.768, *η*^2^ = 0.000; Fig. 3C) and amplitudes (*F* = 0.222, *p* = 0.644, *η*^2^ = 0.000; Fig. 3C). At the middle gait cycle, the foot and shank were significantly delayed in the greater amplitude condition than in the smaller amplitude condition during the acceleration phase (Fig. 4), but they significantly proceeded in the greater amplitude condition during the deceleration phase (Fig. 5).

**Figure 2.**
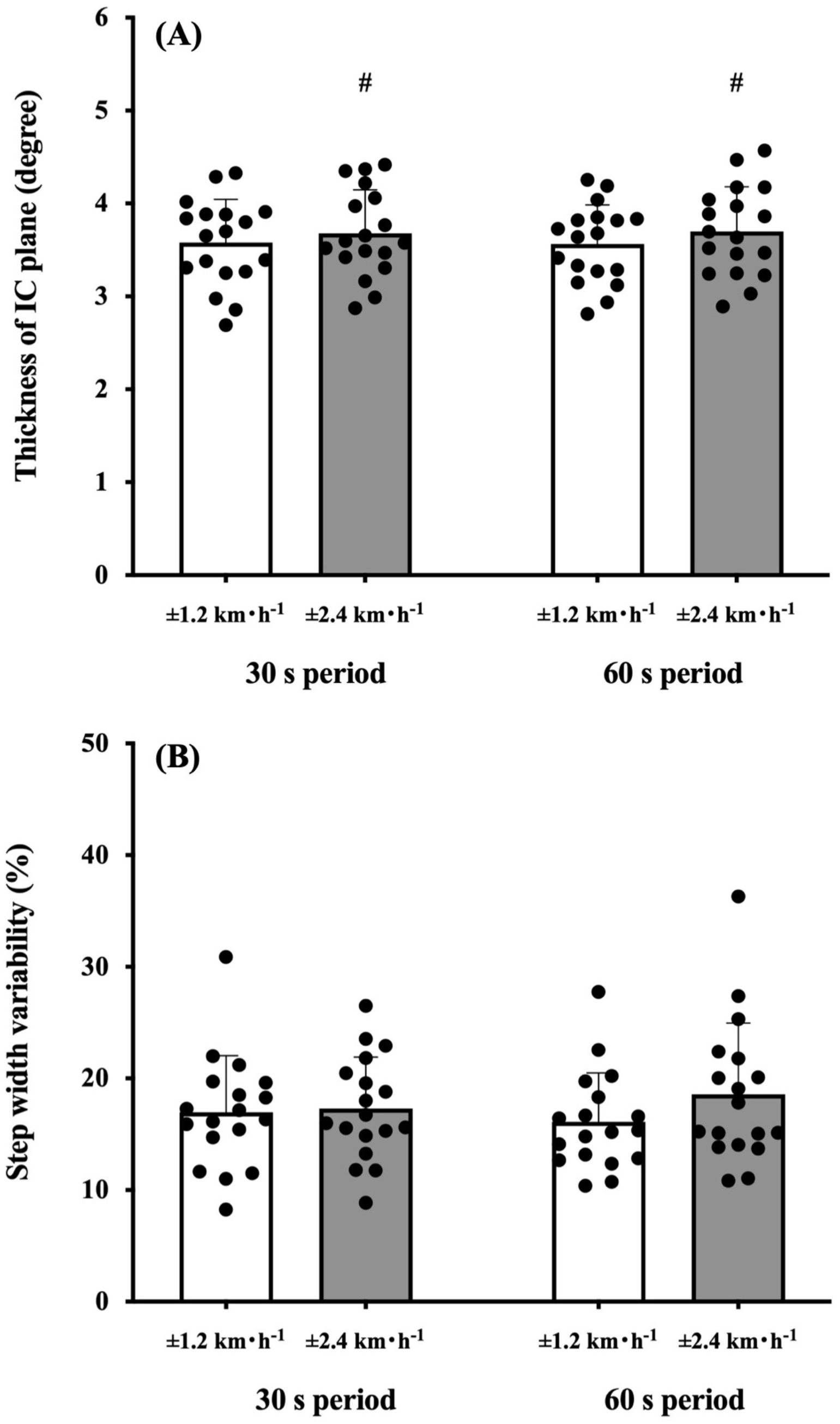
Comparisons of the interlimb coordination (IC) plane thickness and step width variability. **(A)** Participants walked on a treadmill with sinusoidal speed-changing protocols for time periods of 30 s and 60 s periods (left) with amplitudes of ±1.2 km? h^-1^ (white bars) and ±2.4 km? h^-1^ (dark bars), respectively. ±2.4 km? h^-1^ was significantly greater in the IC *plane* thickness. ^#^ *p* < 0.05. **(B)** The coefficient of variance values of the step width variability (CV_SW_; %) were compared between conditions and periods. The CV_SW_ was not significantly different between periods and conditions. Values are presented as means ± standard deviation.

**Figure 3.**
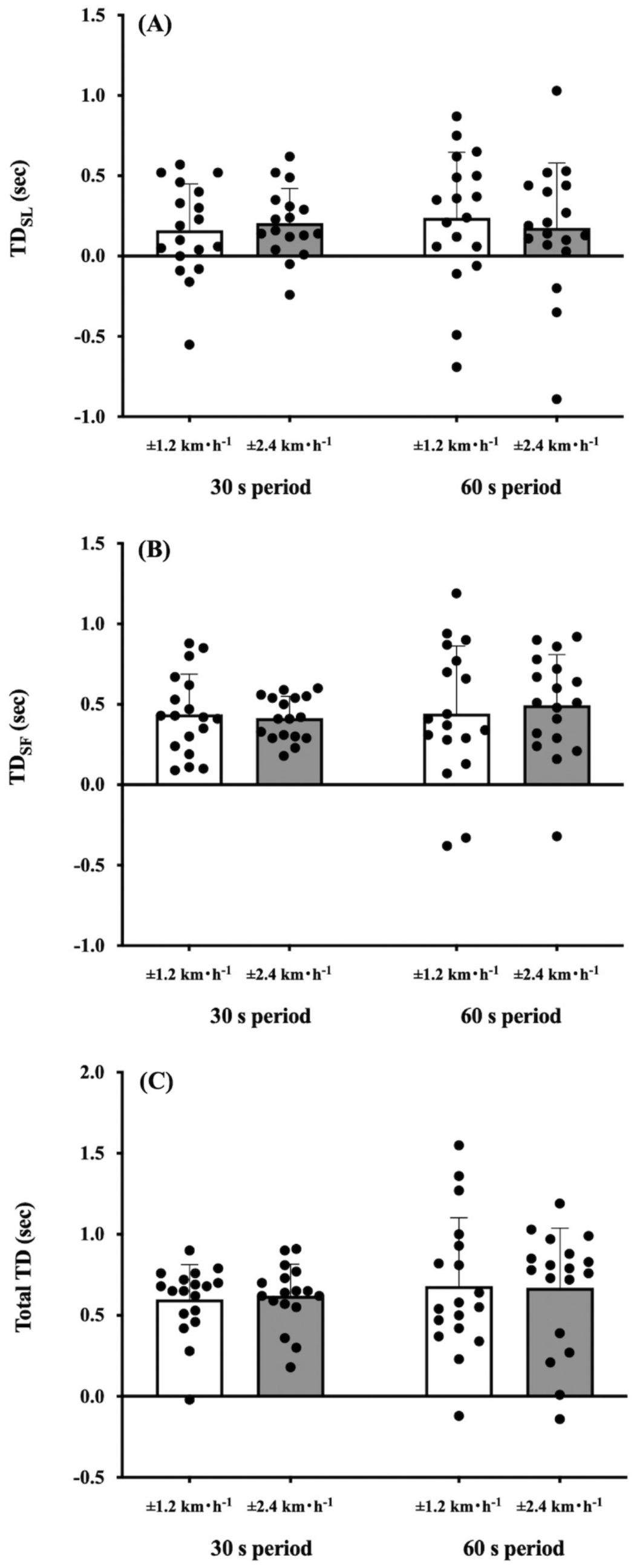
Comparisons of time delay in step variabilities against sinusoidal speed change. **(A)** Time delay (TD) of step length (SL) against sinusoidal speed change. **(B)** TD of step frequency (SF). **(C)** Total TD. No significant differences were found between periods and amplitudes in these parameters. Values are presented as means ± standard deviation.

**Figure 4.**
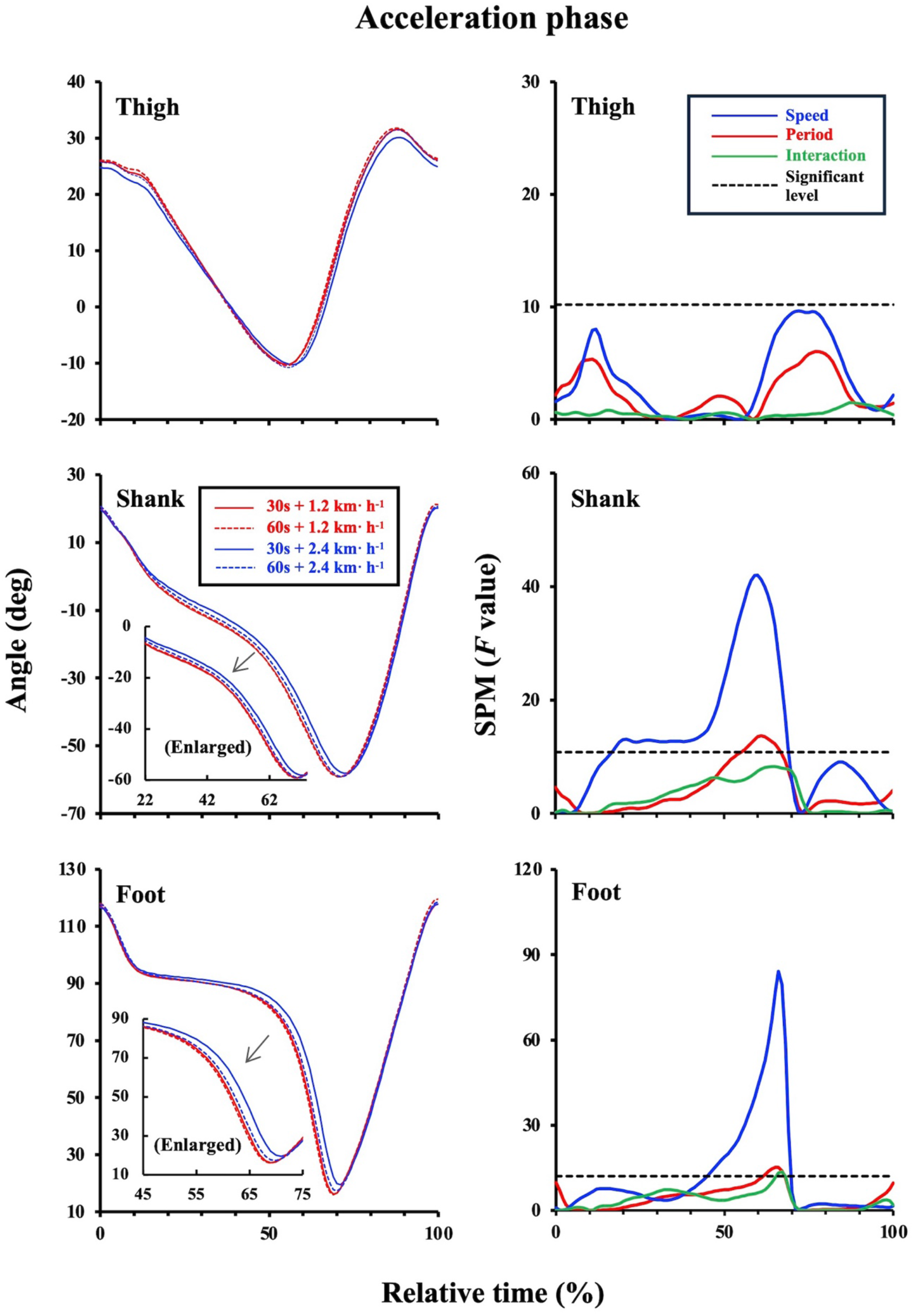
Two-way statistical parametric mapping (SPM) for the relative time series of the lower limbs during the acceleration phase. The left panels represent that the shank and foot were significantly delayed in the ±0.67 m·s^-1^ condition (blue solid and broken lines) than in the ±0.33 m·s^-1^ condition (red solid and broken lines). Enlarged figures were inserted into the left middle panel (*p* < 0.05 at 22%-74%) and left bottom panel (*p* < 0.05 at 45%-75%). The right panels showed statistical results of two-way SPM. The above broken black lines indicate statistical significance (*p* < 0.05). Solid blue, red, and green lines represent the *F* values for the effects of amplitude, period, and interaction, respectively.

**Figure 5.**
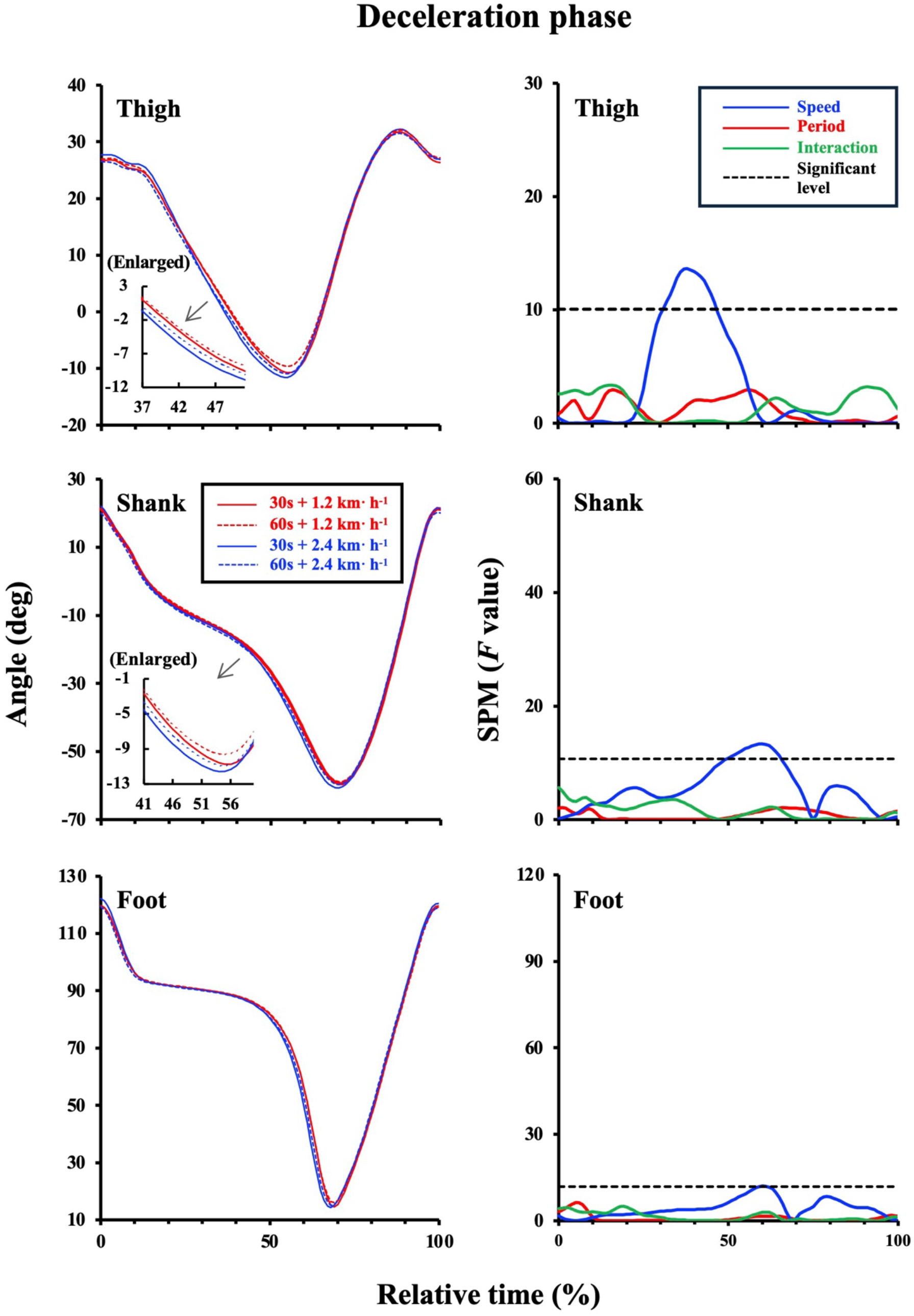
Two-way statistical parametric mapping (SPM) for relative time series of lower limb angles during the deceleration phase. The left panels represent that the thigh and shank were significantly delayed in the ±0.67 m·s^-1^ condition (blue solid and broken lines) than in the ±0.33 m·s^-1^ condition (red solid and broken lines). Enlarged figures were inserted into the upper left (*p* < 0.05 at 37%-51%) and middle left panels (*p* < 0.05 at 41%-60%). The right panels showed statistical results of two-way SPM. Above the broken black lines indicate statistical significance (*p* < 0.05). Solid blue, red, and green lines represent the *F* values for the effects of amplitude, period, and interaction, respectively.

## Discussion

Most of the previous studies have examined the characteristics of the IC *plane* at several steady-state speeds (*Lacquaniti, Grasso & Zago, 1999*; *Hicheur, Terekhov & Berthoz, 2006*; *Ivanenko et al., 2008*; *Bleyenheuft & Detrembleur, 2012*; *Chiu & Chou, 2012*; *Krasovsky et al., 2014*; *Ogaya et al., 2016*; *Wallard et al., 2018*; *Dewolf et al., 2018, 2019*) and demonstrated that gait speed influenced changes in the pattern of the intersegmental coordination of the lower limbs (*Hicheur, Terekhov & Berthoz, 2006*; *Ivanenko et al., 2008*; *Chiu & Chou, 2012*; *Ogaya et al., 2016*; *Dewolf et al., 2018, 2019*). Our recent study revealed that different periods of sinusoidal speed change ranging from 30-s to 120-s did not modify the IC *plane* thickness in young active adults (*Abe et al., 2022*), indicating that anteroposterior gait variability is inherent in each individual. Based on these study backgrounds, we investigated how different amplitudes and periods of sinusoidal speed change influence gait variabilities and/or step variabilities in healthy young adults.

In support of our first hypothesis, the greater the magnitude of the sinusoidal gait speed change, the greater the variability of the IC *plane* thickness (Fig. 2A). The difference of ±0.67 m·s^-1^ and ±0.33 m·s^-1^ condition is the different rate of speed change. That is, the IC *plane* planarity was not necessarily robust if the rate of changing speed increased. In the passive speed changing-condition, appropriate combinations of SL and SF were primarily important to maintain treadmill speed, indicating that efforts to avoid falls are expected to be integrated into step variabilities. Our present study showed that different gait speed changes did not influence TD_SL_ and TD_SF_ (Figs. 3A and B), resulting in a non-significant difference in the total TD among the conditions (Fig. 3C). In addition, the CV_sw_ was not significantly different among the conditions (Fig. 2B), indicating that our second hypothesis was not supported. Rather, our young participants could adjust appropriate combinations of SF and SL during different sinusoidal speed-changing conditions. Previous studies reported that there was a TD between thigh and shank motions even in young adults (*Dewolf et al., 2018, 2019*). Such a TD in the shank-foot coordination may provide greater distortion of the IC *plane* planarity. Thus, some considerations were still necessary because step variabilities are quite large because the coefficient of variance of the total TD was 97.6% (±0.33 m·s^-1^) and 53.3% (±0.67 m·s^-1^) at the 60-s period condition (Fig. 4C), whereas relatively smaller CV_sw_ was found in the ±0.33 m·s^-1^ condition (26.5%) and ±0.67 m·s^-1^ condition (33.5%) at the 60-s period condition (Fig. 2B). Notably, excessive gait variability could be associated with increased fall risks in the elderly population (*Brach et al., 2005*; *Hausdorff, 2007*; *Skiadopoulos et al., 2020*); however, these large variations in the step variabilities may reflect flexible locomotor control ability against passive gait speed changes in healthy young adults.

Further, we compared the relative time series of each limb angle to examine which limbs are attributed to TD_SL_ and/or TD_SF_. The TDs of the thigh-shank and shank-foot decreased as gait speed increased (*Dewolf et al., 2018, 2019*), indicating that followability of the lower extremities was enhanced against treadmill speed particularly at faster gait speed. The shank and foot were significantly delayed in the ±0.67 m·s^-1^ condition than in the ±0.33 m·s^-1^ at the middle gait cycle during the acceleration phase (Fig. 4). Conversely, the thigh and shank significantly proceeded in the ±0.67 m·s^-1^ condition than in the ±0.33 m·s^-1^ condition during the deceleration phase (Fig. 5). That is, more distal limbs were delayed in greater amplitude conditions than in the smaller amplitude conditions at the middle gait cycle during acceleration phase, whereas more proximal limbs proceeded in these conditions during the deceleration phase. Step variabilities refer to the ability of the neuromuscular system to adapt to changing gait conditions (Chiu & Chou, 2012; Ogaya et al., 2016). Consequently, different behaviors of these lower limb segments during the acceleration and/or deceleration phases may be responsible for step variabilities against sinusoidal gait speed changes. If human gait is compared to a bipedal robotics gait, passive walk performed by a bipedal robot intentionally creates an unstable state with a perturbation, and it achieves a stable gait by autonomously controlling the “perturbation state” that causes dynamic imbalance (*McGeer, 1990*; *Taga, 1995*; *Collins et al., 2005*). In that meaning, human gait is quite similar to the passive walk (*McGeer, 1990*). It also characterized by little energy cost (*Donelan et al., 2008*), and this may be related to high efficiency of real human gait (*Abe, Fukuoka & Horiuchi, 2019*; *Ortega & Farley, 2015*). In this study, the distal and proximal limbs behaved differently during the acceleration and deceleration phases in the 60 s cycle at the ±0.67 m/s condition (Fig. 5). In human gait, sinusoidal (continuous) speed change should correspond to the “perturbation”, and slight errors of appropriate SL-SF combination may contribute to achieve successful next new step. The weight balance of each lower limb segment in association with potential “perturbation” characteristics may play an important role in achieving successful gait in the sinusoidal speed changing condition.

### Limitations

We have confirmed that the distal and proximal limbs behaved differently in response to sinusoidal speed changes during the acceleration or deceleration phase. However, our results were obtained from healthy young adults only. Effects of aging (*Abe et al., 2022*; *Horiuchi et al., 2023*) and a lack of exercise habituation (*Abe et al., 2022*) should be considered in the future study.

## Conclusions

The IC thickness was greater in the ±0.67 m·s^-1^ condition than in the ±0.33 m·s^-1^ condition (Fig. 2A). Periods and amplitudes did not affect CV_SW_, TD_SL_, and TD_SF_ (Figs. 2B and 3). The shank and foot were delayed in the ±0.67 m·s^-1^ condition than in the ±0.33 m·s^-1^ condition during the acceleration phase (Fig. 4), whereas the thigh and shank proceeded in the ±0.67 m·s^-1^ condition than in the ±0.33 m·s^-1^ condition during the deceleration phase (Fig. 5). Given these, greater sinusoidal speed change influenced anteroposterior gait variability, but not lateral gait variability, regardless of periods even in healthy young adults. More distal limbs were delayed in greater speed amplitude conditions during the acceleration phase, whereas more proximal limbs proceeded in that condition during the deceleration phase. These different behaviors of the lower limb segments may be attributed to step variabilities.

## Acknowledgements

We specially thank Mr. Akinobu Sakamoto, Mr. Tomokazu Iwatani, Mr. Hiromichi Ikegami, Mr. Takeshi Saito, Mr. Masaru Hashimura, and Mr. Shizuo Takatoh (Takei Scientific Instruments Co., Ltd.) for customizing the treadmill to control the gait speed sinusoidally.

## Authors’ contributions

DA, KM, AS, TS, and MH designed the study. KM, TT, and DA collected and analyzed the data. KM and DA drafted the original manuscript, and then, AS, TT, TS, and MH revised the manuscript. All authors have read and agreed to the final version of the manuscript.

## Funding

This study was financially supported by Grant-in-Aid for Scientific Research from the Japan Society for the Promotion of Science (JP22K11517 to DA, JP20K19623 to KM, and JP21K17613 to AS). Equipment and software installation were partly supported by Grant-in-Aid for Kyushu Sangyo University (KSU) Scientific Research and Encouragement of Scientists (K035124 and R035027 to DA), Special Promotion Research (T018321 to DA, KM, AS, TT, and TS), and Japan Keirin Autorace Foundation (2024M-450 to DA).

## Notes

### Competing Interest Statement

The authors have declared no competing interest.

